# Structural variants and selective sweep foci contribute to insecticide resistance in the *Drosophila melanogaster* genetic reference panel

**DOI:** 10.1101/301937

**Authors:** Paul Battlay, Llewellyn Green, Pontus B. Leblanc, Joshua M. Schmidt, Alexandre Fournier-Level, Charles Robin

**Author notes:** Corresponding author: Charles Robin, Building 184, Parkville, Melbourne Victoria, 3010. Australia.

## Abstract

Patterns of nucleotide polymorphism within populations of *Drosophila melanogaster* suggest that insecticides have been the selective agents driving the strongest recent bouts of positive selection. However, there is a need to explicitly link selective sweep loci to the particular insecticide phenotypes that could plausibly account for the drastic selective responses that are observed in these non-target insects. Here, we screen the Drosophila Genetic Reference Panel with two common insecticides; malathion (an organophosphate) and permethrin (a pyrethroid). Genome wide association studies map ‘survival-on-malathion’ to two of the largest sweeps in the *D. melanogaster* genome; *Ace* and *Cyp6g1*. Malathion survivorship also correlates with lines which have high levels of *Cyp12d1* and *Jheh1* and *Jheh2* transcript abundance. Permethrin phenotypes map to the largest cluster of P450 genes in the Drosophila genome, however in contrast to a selective sweep driven by insecticide use, the derived state seems to be associated with susceptibility. These results underscore previous findings that highlight the importance of structural variation to insecticide phenotypes: *Cyp6g1* exhibits copy number variation and transposable element insertions, *Cyp12d1* is tandemly duplicated, the *Jheh* loci are associated with a *Bari1* transposable element insertion, and a *Cyp6a17* deletion is associated with susceptibility.

## Introduction

Understanding the genetic basis of insecticide resistance is important, not only to inform the implementation of insecticides in agriculture and disease vector control, but also as an evolutionary case-study operating over observable periods of time. Utilizing genome-wide association studies (GWAS) to investigate insecticide resistance provides an unbiased way to identify multiple natural genetic variants associated with a phenotype, while the polymorphism data surrounding associated variants may provide clues to their evolutionary trajectory.

Since its introduction in 2012, the Drosophila Genetic Reference Panel (DGRP; Mackay *et al.* 2012) has proved to be a powerful tool for dissecting the genetic architecture of a range of *Drosophila melanogaster* phenotypes through the implementation of genome-wide association studies (GWAS) on the DGRP’s 205 inbred, sequenced lines. Insecticide-induced mortality has been among these phenotypes (Battlay *et al.* 2016; Deneke *et al.* 2017; Schmidt *et al.* 2017). In 2015, the DGRP’s utility was increased with the introduction of transcriptome data (Huang *et al.* 2015), allowing phenotypes to be tested for association directly with variation in individual transcripts across the *D. melanogaster* transcriptome.

The sequence data generated by the DGRP has also proved to be a valuable resource for the study of population genomics, and has allowed the identification of regions of strong, recent selection in the DGRP’s ancestral population (Garud *et al.* 2015). Two of the most pronounced of these signals, genome wide, come from insecticide resistance loci *Cyp6g1* and *Ace*. Significant selective signals have also been identified around these loci in other *D. melanogaster* populations (Garud and Petrov 2016), and related species (Schlenke and Begun 2004; Signor, New & Nuzhdin 2017; *D. simulans*), as well as by targeted analyses in *D. melanogaster* of selection at *Ace* (Karasov, Messer & Petrov 2010) and *Cyp6g1* (Catania *et al.* 2004; Schmidt *et al.* 2010).

The fact that insecticides appear to have played such an important role in the recent evolutionary history of the DGRP allows us the rare opportunity to study the quantitative genetics of a trait in the process of strong selection. It is unknown, however, what compound or compounds are causing this selection. Although natural variation in both *Ace* and *Cyp6g1* has been demonstrated to confer resistance to various insecticides, attempts to detect these associations in the DGRP, and hence associate the selective sweeps at these loci with a particular compound, have departed from expectations.

Acetylcholineesterase (Ace) is the molecular target of organophosphate insecticides, and four non-synonymous substitutions in the enzyme’s active groove have been demonstrated to reduce the binding capacity of organophosphate insecticides (Menozzi *et al.* 2004). Battlay *et al.* (2016) were, however, unable to detect a significant effect of variation in *Ace* on resistance to the organophosphate azinphos-methyl in DGRP larvae, but instead detected a strong association with alleles that overexpressed *Cyp6g1*, a cytochrome P450 enzyme previously shown to confer metabolic resistance to DDT and imidacloprid when overexpressed (Daborn *et al.* 2001, Daborn *et al.* 2002, Joussen *et al.* 2008, Hoi *et al.* 2014). Although the link between natural alleles which overexpress *Cyp6g1* and resistance to DDT has been demonstrated in a worldwide sample (Catania *et al.* 2004) and Australian populations (Schmidt *et al.* 2010), a similar result was not observed in the DGRP (Schmidt *et al.* 2017).

Aside from the recently reported association between azinphos-methyl resistance and *Cyp6g1* in the DGRP (Battlay *et al.* 2016), previous investigations have mapped organophosphate resistance to a region including *Cyp6g1* (Kikkawa 1961 [parathion]; Pyke *et al.* 2004 [diazinon]). Cross-resistance to the organophosphate malathion was reported at the mapping region on chromosome 2 by Kikkawa (1961), and Ogita’s (1958) mapping of DDT resistance. Le Goff *et al.* (2003) also reported malathion cross-resistance in DDT resistant lab lines (*Hikone*-*R* and *Wisconsin*), both of which showed heightened levels of *Cyp6g1* transcript. Likewise, DDT-resistant *91*-*R* (which carries a resistance allele at the *Cyp6g1* locus [Schmidt *et al.* 2017] and overexpresses the enzyme [Pedra *et al.* 2004]), shows cross resistance to malathion (Misra *et al.* 2013).

In light of this evidence, resistance to organophosphates makes a compelling subject for study in the DGRP. Natural variants with the two strongest signals of selection in the population, *Ace* and *Cyp6g1*, may both confer resistance to these compounds, and the organophospate class of insecticides has been employed widely over a long period of time, giving it opportunity to induce such selective pressures.

Pyrethroids have also been extensively utilized both spatially and temporally in insect control. Natural variation contributing to pyrethroid insecticide class resistance in *D. melanogaster* is less well understood, however they do share the same moleculer target as organochlorides (such as DDT) while being chemically unrelated. Like Ace, resistance-causing mutations in the molecular target of pyrethroids and DDT, the voltage-gated sodium channel, are common in insect pest species (Dong *et al.* 2014). However, orthologous mutations have not been described as natural variants in *D. melanogaster*, although EMS mutagenesis has yielded mutations in *para* (the *D. melanogaster* voltage-gated sodium channel alpha subunit) that cause resistance to DDT and the pyrethroid deltamethrin (Pittendrigh *et al.* 1997). At least one *D. melanogaster* cytochrome P450 gene has bene shown to be involved in pyrethroid biology. *Cyp4e3* is both induced in response to permethrin exposure, and capable of increasing resistance to the insecticide when overexpressed (Terzhaz *et al.* 2015), however, once again natural variation in this gene has not been described, and any contribution to of this locus to pyrethroid resistance in wild populations is yet to be determined.

Organophosphates and pyrethroids are two of the oldest and most widely used insecticide classes in the world today. Here, we investigate the genetic basis of resistance in the DGRP to a representative of each of these classes; the organophosphate, malathion, and the pyrethroid, permethrin. We perform genomic and transcriptomic associations with both male and female adults at multiple doses and incorporate genotyping of structural variation and previously identified signatures of selective sweeps.

## Results

### Genome-wide association studies

Male and female insecticide phenotypes (3, 6, 12 and 24-hour malathion mortality at 1μg/vial, permethrin 3-hour knockdown and permethrin 24-hour mortality at 10 μg/vial) were tested for associations with genomic variants using the DGRP2 pipeline (Huang *et al.* 2014). From the eight malathion phenotypes, we identified 756 unique variants with mixed p-values below 1 × 10^-5^, the arbitrary ‘genome-wide significance’ threshold (fig. 1; fig. S1; table S1). These included 23 nonsynonymous variants in 17 genes, and 95 variants with mixed p-values below 2.66 × 10^-8^, the Bonferroni-corrected significance threshold.

**Figure 1:**
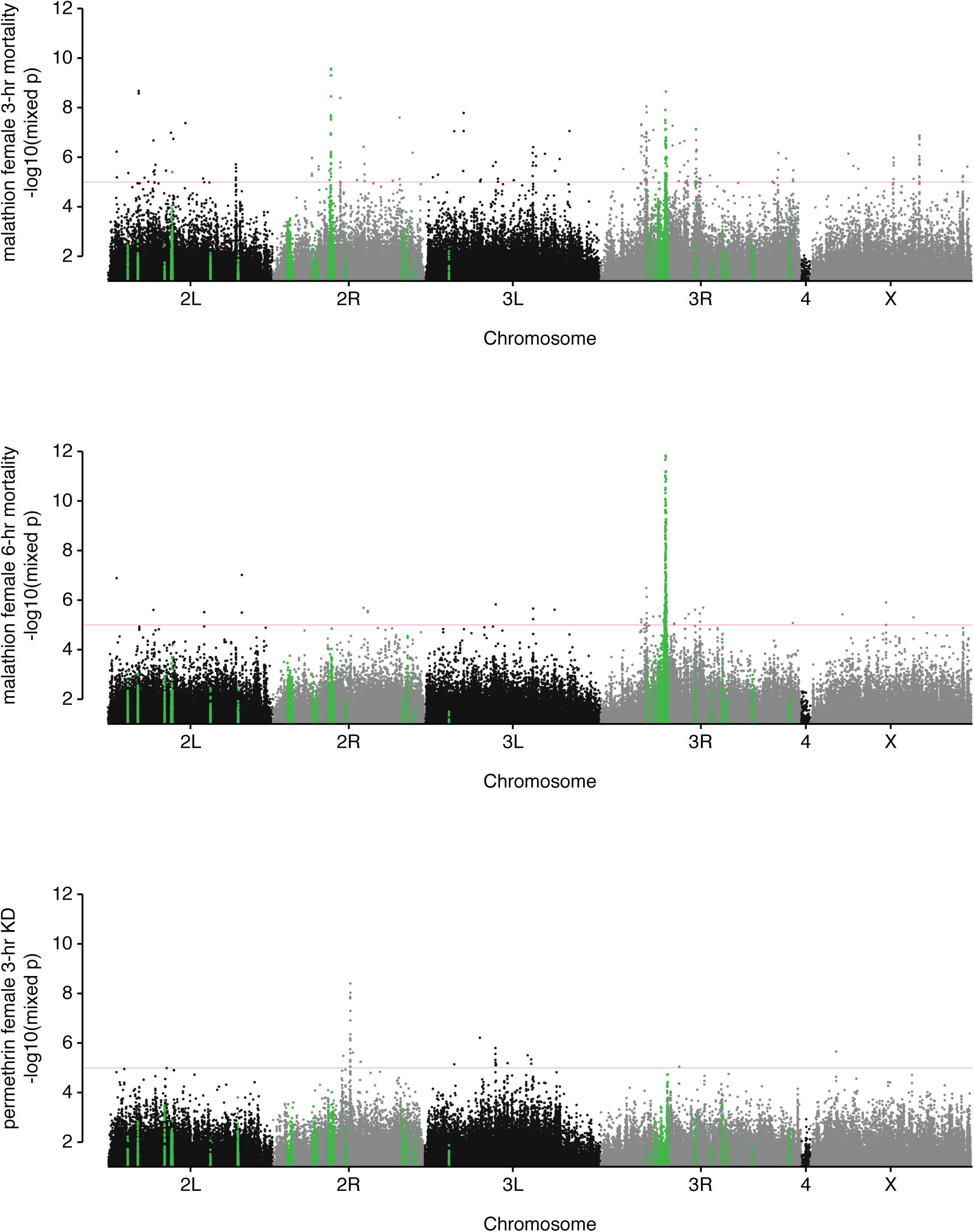
Most significant DGRP associations with resistance to malathion and permethrin. Manhattan plots (mixed p-value vs genomic location) for two malathion phenotypes and one permethrin phenotype, showing strong associations around *Cyp6g1*, *Ace*, and the cytochrome P450 cluster on chromosome 2R. Genome-wide significance threshold (1×10^-5^) indicated in red. Green highlights show variants within DGRP H12 selective sweep windows (Garud *et al.* 2015).

For permethrin, 39 variants were associated with male or female 3-hour knockdown with mixed p-values below 1 × 10^-5^(fig. 1; fig. S1; table S2), four of which achieved Bonferroni-corrected significance (p<2.66 ×10^-8^). All four Bonferroni-significant variants were within the cytochrome P450 cluster on chromosome 2R. For 24-hour mortality, 80 variants were associated with the male or female phenotype with p-values below 1×10^-5^ (fig. S1; table S2, but none reached Bonferroni-corrected significance (p<2.66×10^-8^). Across all permethrin phenotypes, eight nonsynonymous variants in seven genes (*Iyd* (2), *Cyp6a19*, *Cyp6a23*, *CG2444*, *Nha2*, *GlyRS* and *CG6602*) were identified.

### Coincidence with selective sweeps

Variants indicative of haplotypes surrounding *Ace* and *Cyp6g1* were among those strongly associated with malathion phenotypes; these included the three *Ace* resistance substitutions that segregate in the DGRP (I199V [3R_9069054_SNP], G303A [3R_9069408_SNP] and F368Y [3R_9069721_SNP]; Menozzi *et al.* 2004; Battlay *et al.* 2016) and 2R_8072884_INS, the Accord LTR insertion which differentiates the ancestral *Cyp6g1*-*M* allele from resistant *Cyp6g1* alleles in the DGRP (Daborn *et al.* 2002; Schmidt *et al.* 2010; Battlay *et al.* 2016). Selective sweeps involving *Ace* and *Cyp6g1* have previously been described in the DGRP (Garud *et al.* 2015), and our malathion GWAS identified 262 and 29 associated variants from within boundaries of the *Ace* and *Cyp6g1* sweeps respectively. To ascertain whether any other selective sweeps may be associated with our phenotypes, we looked for associated variants within all selective sweep windows identified by Garud *et al.* (2015) in the DGRP. Malathion-associated variants were identified in a total of eight of the DGRP H12 windows (fig. S1; table S3). Only a single permethrin-associated variant was from within a DGRP H12 window, an intergenic SNP 262,773bp from the centre of the *Cyp6g1* sweep (fig. S1; table S3).

### Transcriptome to phenotype associations

Transcriptome to phenotype association tests were performed for malathion and permethrin phenotypes using the DGRP transcriptome data generated by Huang *et al.* (2015). For malathion, 36 transcripts below an arbitrary p-value of 1 × 10^-4^ were identified, two of which (*Cyp6g1* and *Sf3a1*) were below the Bonferroni-corrected significance threshold (2.76×10^-6^; table S5). Activation of the CncC/Keap1 pathway has previously been shown to confer malathion resistance in *D. melanogaster* (Misra *et al.* 2011). Of our top 36 transcriptome candidates, 12 were among those found by Misra *et al.* (2011) to have their expression altered by two-fold or more by ectopic expression of *CncC*. These include *Cyp12d1*-*p*, a cytochrome P450 enzyme that confers resistance to DDT and dicyclanil when overexpressed (Daborn *et al.* 2007), and *Jheh1* and *Jheh2*, shown by Guio *et al.* (2014) to increase resistance to malathion when induced. In the case of permethrin, four transcripts below an arbitrary p-value of 1×10^-4^ were identified, none of which were below the Bonferroni-corrected significance threshold (table S4).

We also interrogated datasets generated by Huang *et al.* (2015) for insecticide-associated eQTLs whose target transcript was also associated with resistance to the same insecticide. As expected, malathion-associated variants within the *Cyp6g1* sweep boundaries were identified and can be considered ‘cis’ eQTLs for *Cyp6g1.* Likewise, permethrin-associated variants in *Cyp6a23* were eQTLs of *Cyp6a17* in both sexes. While *Cyp6a17* transcript does not reach significance at 1×10^-4^ in any of our transcriptome to phenotype association tests, it has a high rank in all phenotypes (male 3-hour knockdown rank=26, r^2^=0.060; female 3-hour knockdown rank=4, r^2^=0.064; male 24-hour mortality rank=26, r^2^=0.057; female 24-hour mortality rank=26, r^2^=0.066).

### Structural variation in candidate genes

Structural variation among DGRP lines has previously been reported for association study candidates *Cyp6g1*, *Cyp12d1*-*p*, *Jheh1/Jheh2* and *Cyp6a23* (Zichner *et al.* 2013; Good *et al.* 2014; Guio *et al.* 2014; fig. 2). Of these structural variants, only *Cyp6g1* is directly called in DGRP genotype data (2R_8072884_INS encodes *Accord* transposable element insertion status, present in all derived *Cyp6g1* alleles). Therefore, presence of structural variants was directly tested for association against the relevant insecticide phenotype, and significant associations were found with *Cyp6g1* derived alleles in the case of malathion, and the *Cyp6a17* deletion allele in the case of permethrin (two tailed t-test assuming unequal variances, p<0.05; table S5).

**Figure 2:**
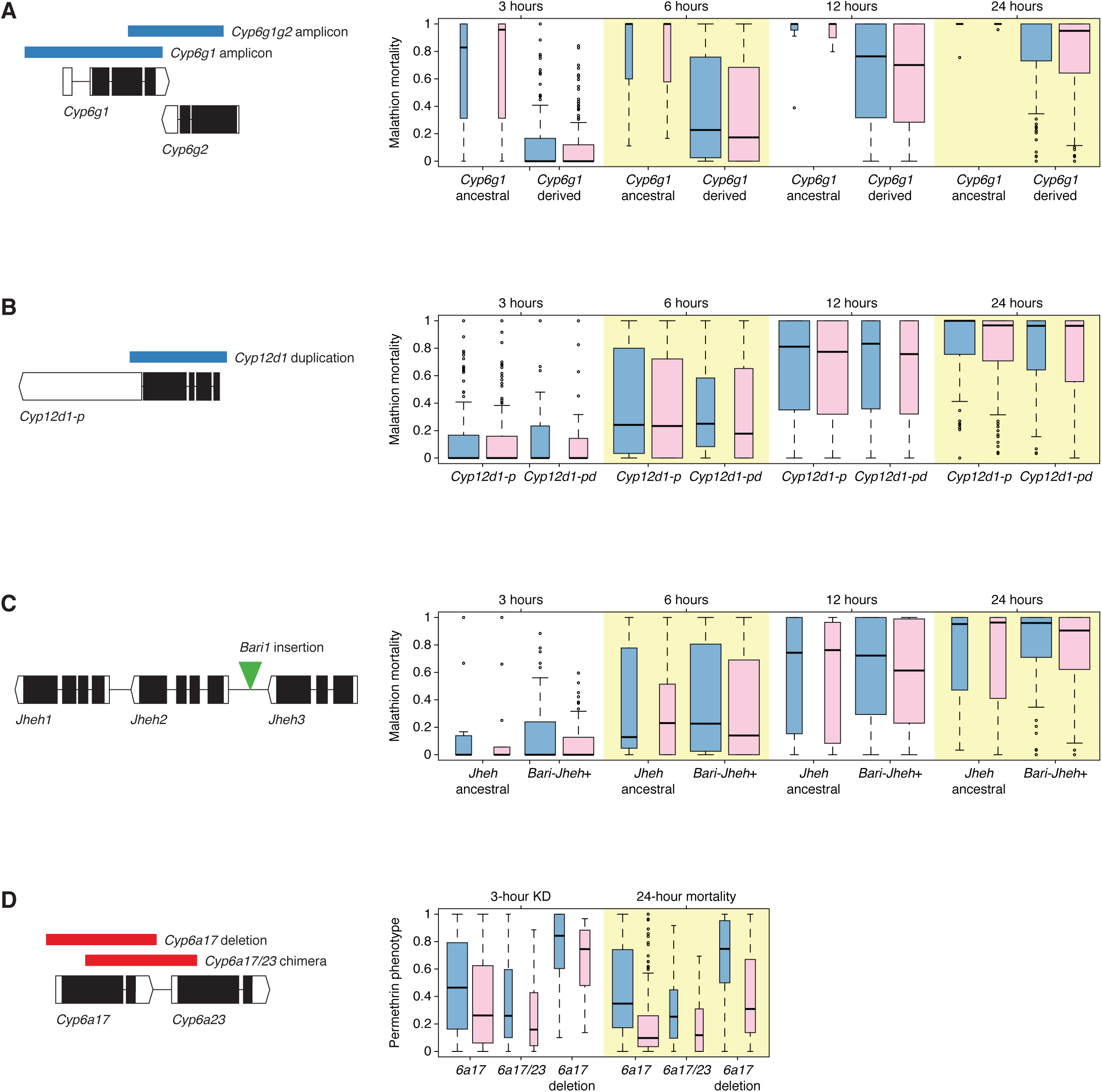
Structural variation in candidate insecticide resistance genes. DGRP structural variation in candidate insecticide resistance genes **(A)** *Cyp6g1*, **(B)** *Cyp12d1*, **(C)** *Jheh1* and *Jheh2*, and **(D)** *Cyp6a17* and *Cyp6a23* (Good *et al.* 2014; González *et al.* 2008). Box plots show phenotype distributions among DGRP lines at all timepoints, grouped by structural variant allele. Blue represents males and pink represents females. Mean phenotypes for *Cyp6g1* derived alleles and *Cyp6a17* deletion alleles are significantly different than reference alleles at in all relevant phenotypes (table S5).

Discordant paired-end read mapping over the *Cyp6g1* and *Cyp6g2* loci shows that the two gene amplification events described in Schmidt *et al.* (2010) are present in the *Cyp6g1*-*AA* and *Cyp6g1*-*BA* alleles among DGRP lines. However, read depth across the region suggests substantial variation in copy number of both these amplicons, implying 1-5 copies of the *Cyp6g1* amplicon, and 1-10 copies of the partial *Cyp6g1g2* amplicon, which are correlated with transcript levels of both *Cyp6g1* and *Cyp6g2* (fig. S2).

### Knockout of *Cyp6g1* increases susceptibility to three organophosphate insecticides

Natural variation at *Cyp6g1* contributes to resistance to a range of insecticides including DDT (Schmidt *et al.* 2010), azinphos-methyl (Battlay *et al.* 2016) and imidacloprid (Deneke *et al.* 2017) and also ranks at the top of our genomic and transcriptomic association tests with malathion phenotypes. We verified *Cyp6g1* involvement in malathion resistance using RAL_517-*Cyp6g1*-KO, a DGRP line in which the natural resistance allele *Cyp6g1*-*BA* is knocked out (Deneke *et al.* 2017). We observed a decrease in LD_50_ of approximately two thirds in both male and female RAL_517-*Cyp6g1*-KO flies when compared to unmodified RAL_517 flies (fig. 3).

**Figure 3.**
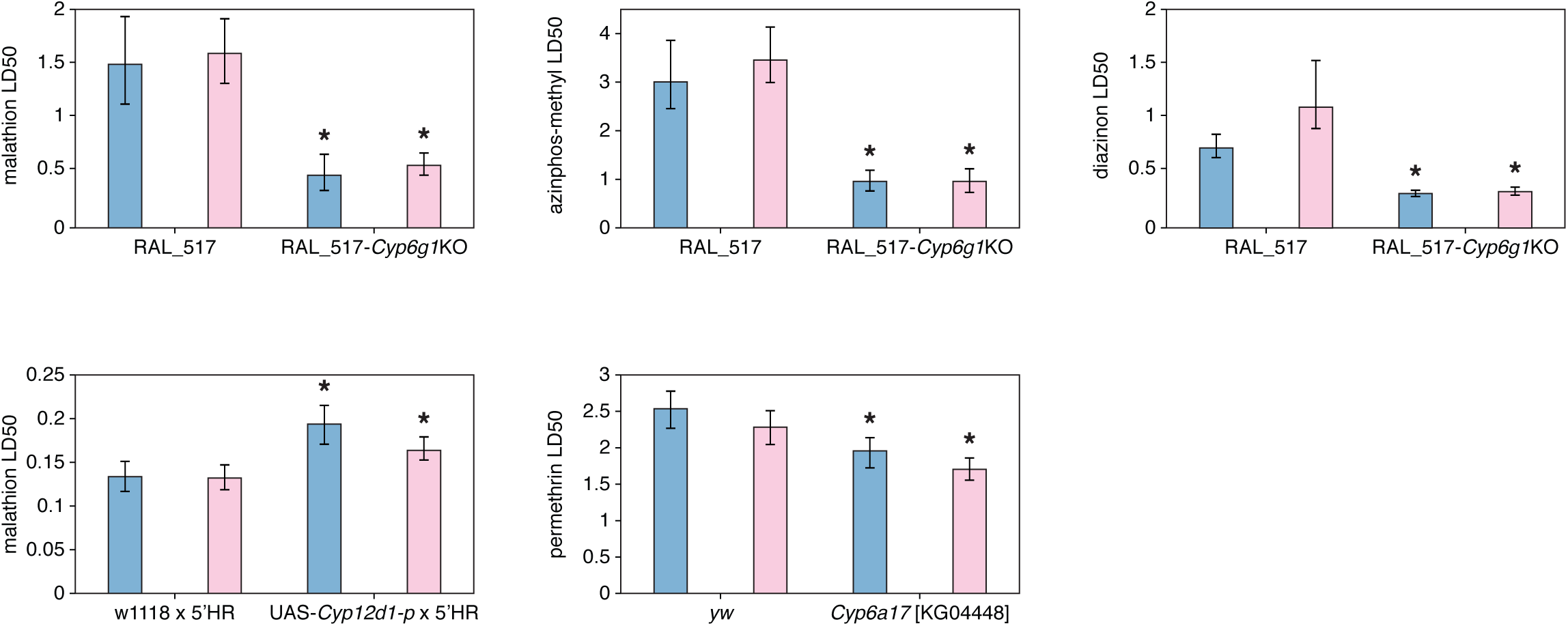
Functional validation of insecticide resistance candidates. Insecticide LD_50_s for both male and female adults. Error bars represent 95% confidence interval. Blue represents males and pink represents females. CRISPR knockout of *Cyp6g1* in the DGRP line RAL_517 background significantly increases susceptibility to organophosphate insecticides malathion, azinphos-methyl and diazinon. Transgenic overexpression of *Cyp12d1*-*p* with the *6g1*HR-GAL4 driver significantly increases resistance to malathion. Gene Disruption Project line *Cyp6a17^KG04448^* shows increased permethrin susceptibility.

Battlay *et al.* (2016) previously demonstrated that *Cyp6g1* overexpression, both transgenic (using GAL4-UAS overexpression) and among DGRP lines, was associated with resistance in larvae to the organophosphate azinphos-methyl. Pyke *et al.* (2004) mapped resistance to another organophosphate, diazinon, in an Australian natural population of *D. melanogaster* to a region containing *Cyp6g1*. Here we also present toxicological assays of RAL_517-*Cyp6g1*-KO and RAL_517 that demonstrate that the removal of the natural *Cyp6g1*-*BA* resistance allele from RAL_517 significantly reduces resistance to azinphos-methyl and diazinon in male and female adults (fig. 3).

### *Cyp12d1*-*p* overexpression increases malathion resistance

Increased expression of *Cyp12d1* has previously been linked with resistance to the insecticides DDT and dicyclanil (Pedra *et al.* 2004; Daborn *et al.* 2007; Gellatly *et al.* 2015). In this study, we detected an association between male mortality at 24 hours and transcript level of *Cyp12d1*-*p* (adjusted r^2^=0.12; p=4.80×10^-6^), one of the two copies of *Cyp12d1* present in the *y; cn bw sp;* genome. Flies overexpressing *Cyp12d1*-*p* had significantly higher LD_50_s (~30% and ~20% increases for males and females respectively) than control crosses (fig. 3).

### *Cyp6a17* disruption increases permethrin susceptibility

Investigation of variation in the cytochrome P450 cluster on chromosome 2R shows that variants associated with permethrin phenotypes are in linkage disequilibrium with deletions of *Cyp6a17*. To test the hypothesis that *Cyp6a17* contributes to permethrin resistance, we obtained a Gene Disruption Project line (Bellen *et al.* 2004), *Cyp6a17^KG04448^*, in which a P-element construct had been inserted into the coding region of *Cyp6a17*, early in the first exon. We found significantly reduced LD_50_s in both males and females from this line when compared to control flies (fig. 3).

## Discussion

In this study, the DGRP was assayed for resistance to malathion and permethrin, representatives of two of the most important insecticide classes, organophosphates and pyrethroids. Sexes were phenotyped separately and scored at multiple time points, increasing the resolution with which variants contributing to resistance could be identified. This is evident in the variation in the strength of association across time points, even for the top candidates *Cyp6g1*, *Ace*, and *Cyp6a17/Cyp6a23* (Fig. S1).

Evidence suggests that insecticides have played a large role in recent selection in *D. melanogaster*. Here, we find insecticide-associated variants in eight of the 25 H12 selective sweep windows identified in the DGRP by Garud *et al.* (2015). However only the windows containing *Cyp6g1* and *Ace* had multiple insecticide-associated variants within them. The other sweep windows may associate with these insecticides via more subtle soft sweeps, be more strongly associated with other insecticides, or be driven by other selective agents. Nevertheless there is strong evidence to support the hypothesis that both *Cyp6g1* and *Ace* are the targets of selection in the population ancestral to the DGRP and other populations (Catania *et al.* 2004; Schmidt *et al.* 2010; Karasov *et al.* 2010; Garud *et al.* 2015; Garud & Petrov 2016), and the link to malathion resistance is now compelling.

Of the four substitutions in *D. melanogaster Ace* known to confer enzymatic insensitivity and hence resistance to organophosphate and carbamate insecticides (Menozzi *et al.* 2004), the three most common are present in DGRP lines (I199V, G303A and F368Y). Each of these three nonsynonymous sites independently achieved Bonferroni-significant associations with male and female malathion mortality at 6 and 12 hour timepoints. Two resistant *Ace* substitution haplotypes exist among DGRP lines; *Ace-*VGF (one substitution) and *Ace*-VAY (three substitutions), and enzymes carrying these substitutions have inhibitory constants of 6.4 and 32 to malaoxon (the activated form of malathion) respectively (Menozzi *et al.* 2004). We find that DGRP population mean malathion mortalities for each of the three haplotypes in keeping with these relationships, suggesting that the previously characterized role of *Ace* resistance substitutions explains the strong malathion associations detected within *Ace* and the surrounding haplotype. Previously, a DGRP GWAS of resistance to the organophosphate azinphos-methyl did not detect strong associations with these alleles (Battlay *et al.* 2016). This is likely due to the lower inhibitory constants of these alleles to azinphos-methyl than malathion; *Ace-* VGF actually reduces the inhibitory constant to 0.92, and *Ace*-VAY only increases it to 4.8 (Menozzi *et al.* 2004).

Derived alleles of *Cyp6g1* are associated, through both genomic and transcriptomic variation, with malathion resistance. This study adds to the mounting evidence that *Cyp6g1* overexpression in wild populations confers resistance to a range of organophosphate insecticides (Kikkawa 1961; Pyke *et al.* 2004; Battlay *et al.* 2016), and that organophosphate selection may be a more likely explanation than DDT for the sweep observed at the *Cyp6g1* locus (Schmidt *et al.* 2017). Moreover, we find that the DGRP harbors greater allelic diversity at *Cyp6g1* than had previously been described at the locus, and that these additional structural variants, along with those previously characterized, are correlated with differences in transcription of *Cyp6g1* and downstream *Cyp6g2.*

This work also implicates cytochrome P450s in resistance to permethrin. DGRP variants most strongly associated with permethrin map to a region on chromosome 2R containing nine P450 genes, with peaks over *Cyp6a23* and *Cyp317a1*. Structural variation in the DGRP has previously been described involving *Cyp6a23* and its tandem paralog *Cyp6a17* (Zichner *et al.* 2013; Good *et al.* 2014). Two deletions segregate in this region, one creates a single chimeric gene comprised of *Cyp6a17* and *Cyp6a23* sequence. In the other, *Cyp6a17* is deleted, save for a small section which exists as a gene conversion in the otherwise intact *Cyp6a23*. Due to the homology between these genes, this gene conversion introduces only four nucleotide changes and a single non-synonymous substitution in *Cyp6a23* (Good *et al.* 2014). Among DGRP lines, it is this deletion of *Cyp6a17* that is associated with increased susceptibility to permethrin (fig. 2D). This is congruent with the finding that when *Cyp6a17* is disrupted in *Cyp6a17^KG04448^*, it leads to relative susceptibility (fig. 3). It is worth noting that the susceptible allele appears to be the derived state, as the duplication leading to the original divergence *Cyp6a17* and *Cyp6a23* seems to be at least as old as the divergence between *D. melanogaster* and *D. ananassae* (Good *et al.* 2014).

Interestingly, *Cyp6a17* is ranks fourth among *D. melanogaster* P450s (after *Cyp6a13*, *Cyp6a2*, and *Cyp6a14*) in similarity to malaria vector *Anopheles funestus CYP6P9a* and *CYP6P9b*. Naturally occurring duplications of each of these genes are associated with pyrethroid resistance in *A. funestus* (Wondji *et al.* 2009), and selective sweeps have been described at *CYP6P9a* in response to pyrethroid-based malaria interventions (Barnes *et al.* 2017).

The transcription level of another P450, *Cyp12d1*-*p*, was associated with male 24-hour mortality, and transgenic overexpression of the gene confers malathion resistance in both male and female adults (fig. 3). Interestingly, we do not observe a strong association at either the genomic level (table S5) or the transcriptome level (table S4) with *Cyp12d1*-*d*, present as a duplicated paralog in 24% of DGRP lines. Coding differences (three amino acids differentiate *Cyp12d1*-*p* and *Cyp12d1*-*d* in the reference genome) or expression patterns between the genes may explain this observation. Alternatively, *Cyp12d1*-*p*’s significance may be inflated by its strong correlation with a group of co-regulated genes that are known to be induced by oxidative stress (Misra *et al.* 2011). We found that nine top malathion-associated transcripts are among those known to be regulated by CncC (Misra *et al.* 2011), and they are clustered together in Huang *et al.* (2015)’s genetically correlated transcription modules (table S4).

Two more CncC-activated transcripts associated with malathion phenotypes are *Jheh1* and *Jheh2*. Guio *et al.* (2014) demonstrated that the insertion of *Bari1* upstream of *Jheh1* and *Jheh2* increases the inducibility of these genes in response to oxidative stress, which results in an increased resistance to malathion. In this study, we found associations between constitutive transcript levels of *Jheh1* and *Jheh2* and male malathion mortality at 24 hours (adjusted r^2^=0.10; p=4.52×10^-5^) and 3 hours (adjusted r^2^=0.10; p=4.62×10^-5^) respectively. However, we did not find that the presence of the *Bari*-*Jheh* insertion was significantly associated with malathion mortality at any of our four timepoints, for either sex (fig. 2; table S5). This is presumably due to differences in the exposure times between our study (up to 24 hours) and the work in Guio *et al.* (2014; up to 214 hours). It makes sense that in our more acute assays, the baseline expression level of expression of *Jheh1* and *Jheh2* would be important, whereas over longer assay periods induction capacity would play a more important role.

An important question in DGRP insecticide resistance studies is the applicability of findings in this subset of variation to populations worldwide. *Ace* and *Cyp6g1* resistance alleles have been identified in a range of populations, as have the footprints of their selection (Catania *et al.* 2004; Karasov, Messer & Petrov 2010; Schmidt *et al.* 2010; Garud & Petrov 2016). This is further reinforced by interrogation of the Drosophila Genome Nexus (DGN; Lack *et al.* 2016), which reveals *Ace* and *Cyp6g1* allele frequencies comparable to the DGRP in many populations around the world (table S7). The *Bari*-*Jheh* allele and *Cyp12d1* duplication have also been described in populations outside the DGRP (González, Macpherson & Petrov 2009; Kolaczkowski *et al.* 2011).

In this investigation of insecticide resistance in the DGRP using an organophosphate and a pyrethroid insecticide we saw stark differences in the genes involved as well as the evidence for a selective response to the compounds. Malathion top candidates correlate with peaks of selection in the DGRP population, suggesting that malathion or a related compound imposed a strong selective pressure on the population ancestral to the DGRP lines. Conversely, *Cyp6a17*, our top permethrin candidate, does not lie within a selective sweep window, nor would we expect the allele described to be the target of positive insecticide-based selection, given it increases susceptibility to permethrin. However, common threads running through this work and previous investigations of insecticide resistance in the DGRP suggest that complex structural variation and high allelic diversity, along with selective sweep signatures, are common in genes contributing to resistance.

## Materials and Methods

### Fly lines

DGRP lines were generated by Mackay *et al.* (2012) and obtained from the Bloomington Drosophila stock center in Indiana. The *6g1*HR-GAL4 driver line was generated by Chung *et al.* (2007). The RAL_517 *Cyp6g1*-KO line was generated by Deneke *et al.* (2017). The UAS-*Cyp12d1*-*p* line was generated by Daborn *et al.* (2007). *Cyp6a17^KG04448^* flies were generated by Bellen *et al.* (2011) and were obtained from the Bloomington Drosophila Stock Center. All fly stocks were maintained at 25°C on rich medium containing, maltose (46g/L), dextrose (75g/L), yeast (35g/L), soy flour (20g/L), maize meal (73g/L), agar (6g/L), acid mix (14ml/L), and tegosept (16ml/L). The acid mix solution was made up of orthophosphoric acid (42ml/L), and propionic acid (412ml/L), while the tegosept solution was 50g tegosept dissolved in 950 ml of 95% EtOH.

### Insect bioassays

Adult flies for bioassays were anaesthetised with CO_2_ at 0-24 hours after eclosion and sorted by sex into holding vials containing rich media, where they were kept for 3-4 days, resulting in 3-5 day old adults for use in bioassays. Assays commenced between 11am and 12pm. 20mL glass scintillation vials were treated with 500μl of acetone/insecticide solution at the required concentration and rolled using a hot dog warmer (heat off) until the acetone had evaporated. ~7 flies were transferred to each vial without the use of anesthesia, and cotton wool moistened with 10% sucrose solution was used to stopper the scintillation vials. DGRP lines were screened at a single dose (1μg/vial for malathion, 10μg/vial for permethrin). Malathion-treated flies were scored for mortality at 3, 6, 12 and 24 hours, permethrin-treated flies were scored for knockdown at 3 hours and mortality at 24 hours. Transgenic lines were screened at multiple doses and scored for mortality at 24 hours. A minimum of three biological replicates were performed for each sex of each DGRP line, and for each sex at each dose for transgenic lines.

### Calculation of LD_50_

For transgenic lines, linear models were fitted to dose-mortality data on a log-probit scale using ‘glm’ in the R statistical package (R Core Team 2013) and scripts from Johnson *et al.* (2013). 50% lethal dose (LD_50_) values and 95% confidence intervals were calculated using Fieller’s method from fitted linear models (Finney 1971).

### Genome-wide association studies

Phenotype files for 170 DGRP lines, consisting of mean mortality data for both males and females, were generated for all phenotypes and were submitted to the Mackay lab DGRP2 GWAS pipeline (http://dgrp.gnets.ncsu.edu/data.html; Huang *et al.* 2014).

### Transcriptome to phenotype associations

Transcriptome data for 1-3 day old adult flies from 185 DGRP lines were recovered from the DGRP website (http://dgrp.gnets.ncsu.edu/data.html; Huang *et al.* 2015). Mean transcription level was calculated for each gene in each sex from two biological replicates, to give a mean level for each of the 18,140 transcripts measured by Huang *et al.* (2015) in each DGRP line, for both males and females. A linear model was fit between mean transcription level of each gene measured by Huang *et al.* (2015) for the relevant sex, and mean malathion mortality at 3, 6, 12 and 24 hours for both males and females.

### Genotyping of structural variation

.bam files containing alignments of DGRP line sequences from Illumina platforms to the *y; cn bw sp;* reference genome were recovered from the Baylor College of Medicine website (https://www.hgsc.bcm.edu/content/dgrp-lines; Mackay *et al.* 2012). Local alignments at candidate loci were visualized with IGV 2.0 software (Robinson *et al.* 2011) to manually score structural variation. *Cyp6g1*, *Cyp6a17/23*, and *Cyp12d1* structural variants were previously genotyped in Good *et al.* (2014).

Genotyping of the *Bari1* insertion presence upstream of *Jheh1* and *Jheh2* genes was provided by Josefa González derived from diagnostic PCR (33 lines; González, Macpherson & Petrov 2009) and T-lex software (119 lines; Fiston-Lavier *et al.* 2011). These datasets were supplemented with our own manual calling of the insertion using IGV 2.0 software (167 lines; Thorvaldsdóttir *et al.* 2013). This resulted in 80 DGRP lines with high confidence (at least two concurrent calls) *Bari*-*Jheh* genotypes that also had matching transcriptome and malathion phenotype data, which were used for further analysis.

Amplification events involving *Cyp6g1* and *Cyp6g2* were inferred from local read depth in DGRP .bam files. Read depth at each nucleotide position under consideration were recovered using the Genome Analysis Toolkit ‘DepthOfCoverage’ utility (McKenna *et al.* 2010). Regions interrogated were non-overlapping portions of the *Cyp6g1* amplicon (2R:8072727-8074976), the *Cyp6g1g2* amplicon (2R:8075688-8077656), and a control region of similar size just upstream of *Cyp6g1* which does not exhibit structural variation in the DGRP (2R:8070657-8072656). Mean read depth for each amplicon was calculated for each DGRP line, and normalized to mean read depth of the control region.

### Transgenic overexpression

Candidate metabolic resistance genes were overexpressed using the GAL4/UAS system (Brand and Perrimon 1993) and the *6g1*HR-GAL4 driver described by Chung *et al.* (2007). *6g1*HR-GAL4 virgin females, in which GAL4 is regulated by *Cyp6g1* upstream sequence originating from Hikone-R line flies, were crossed to males carrying an additional copy of candidate gene coding region under control of a UAS promoter. UAS-null is used as a control for UAS-*Cyp6g1* and UAS-*Cyp6g2* (Deneke *et al.* 2017), and w1118 is used as a control for UAS-*Cyp12d1*-*p* and UAS-*Cyp12d1*-*d* (Daborn *et al.* 2007).

### Frequencies of *Ace* and *Cyp6g1* in the Drosophila Genome Nexus

.fasta files from the Drosophila Genome Nexus release 1.1 (Lack *et al.* 2016) were downloaded from http://www.johnpool.net/genomes.html. The provided scripts were used to mask data for identity by descent and population admixture. Variants were retrieved from the genomes using the provided dataslice.pl script. In the case of *Cyp6g1*, we used 2R:8072837, a SNP in complete linkage disequilibrium with derived alleles of *Cyp6g1* in the DGRP, as a marker for derived *Cyp6g1* alleles in the DGN data.

Most of the fly stocks are available at stock centres. All other material and any data that is not presented in the manuscript, is available on request.

## Acknowledgements

Josefa González, Phil Batterham/Batterham lab (stocks), Trudy Mackay, Owain Edwards, NECTAR computational services

## Figure legends

**Figure S1:**
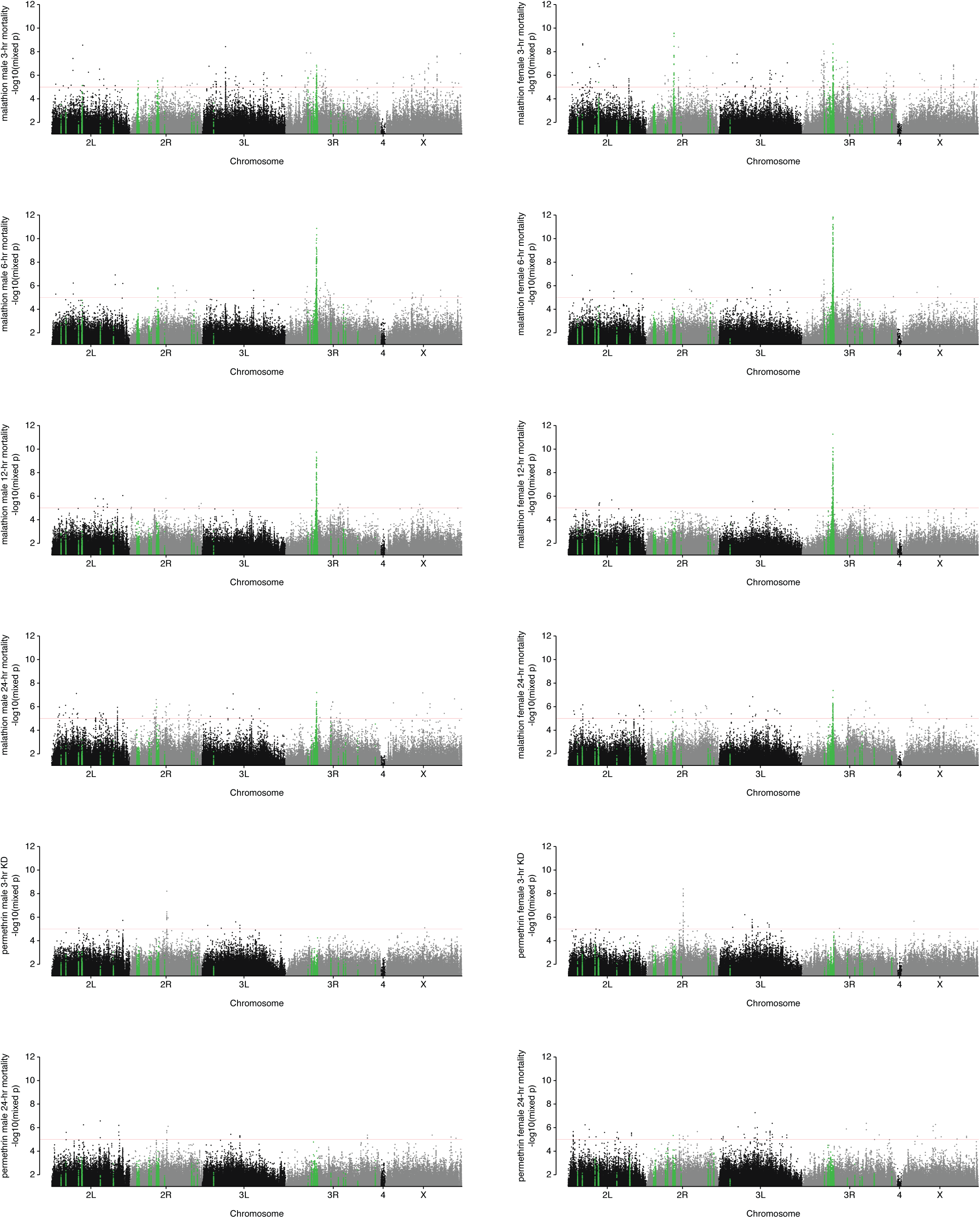
DGRP genomic variants associated with resistance to malathion and permethrin. Manhattan plots (mixed p-value vs genomic location) for eight malathion phenotypes and four permethrin phenotypes. Genome-wide significance threshold (1×10^-5^) indicated in red. Green highlights show variants within DGRP H12 selective sweep windows (Garud *et al.* 2015).

**Figure S2:**
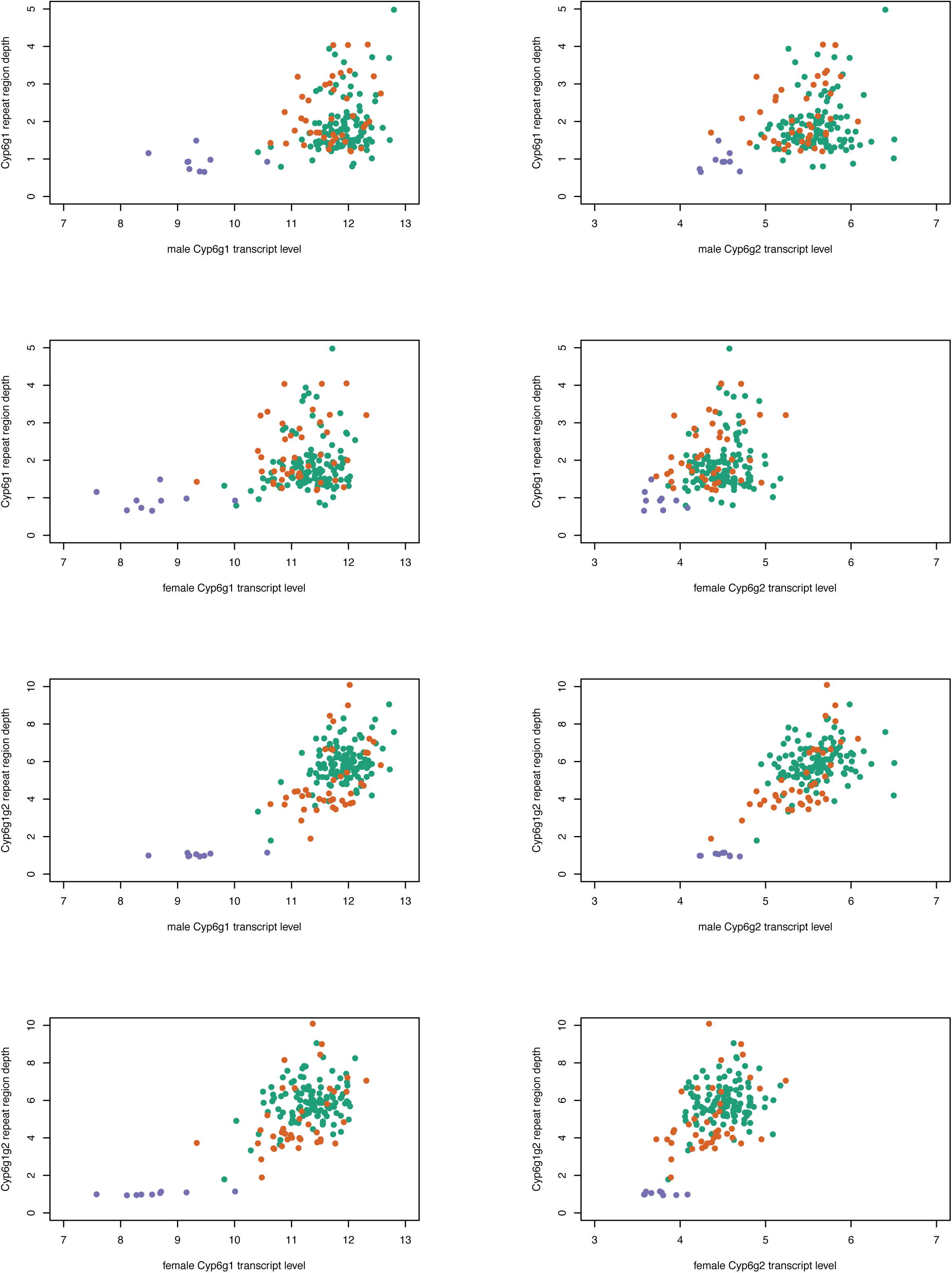
Gene amplification events at *Cyp6g1* and *Cyp6g2* is correlated with transcript levels of both genes. Normalized read depth of the two repeat regions (refer to figure 3) plotted against transcript levels of each gene in male and female adults. Colours represent transposable element insertion alleles (Schmidt 2010; M: purple, AA: Orange, BA: Green)

**Table S1: Variants associated (mixed p<1×10^-5^) with malathion phenotypes in the DGRP.** ^⋆^Denotes variants or groups of variants which include a nonsynonymous site ^⋆⋆^ Yellow highlight denotes group includes a Bonferroni-significant (mixed p<2.66×10^-8^) variant

**Table S2: Variants associated (mixed p<1×10^-5^) with permethrin 3-hour knockdown in male or female DGRP lines.** ^⋆^Denotes variants or groups of variants which include a nonsynonymous site ^⋆⋆^Yellow highlight denotes group includes a Bonferroni-significant (mixed p<2.65×10^-8^) variant

**Table S3: Insecticide phenotype-associated (mixed p<1×10^-5^) DGRP variants within H12 selective sweep windows from the DGRP (Garud *et al.* 2015).**

**Table S4: Transcriptome to phenotype association candidates (p<1×10^-4^) for all malathion and permethrin phenotypes.** ^⋆^ Genetically correlated transcriptional module (Huang *et al.* 2015)

**Table S5: Results of t-tests comparing DGRP phenotype means partitioned by structural variants in candidate genes.**

**Table S6. Insecticide phenotype-associated DGRP variants annotated to genes, and differentiated between *91*-*R* and *91*-*C*, with the resistance-associated allele present in *91*-*R*.** ^⋆^Denotes variants or groups of variants which include a nonsynonymous site

**Table S7: Allele frequencies of *Ace* and *Cyp6g1* resistance alleles in Drosophila Genome Nexus populations (Lack *et al.* 2016)**. ^⋆^ 2R:8072837, a SNP in complete linkage disequilibrium with derived alleles of *Cyp6g1* in the DGRP was used as a marker for derived *Cyp6g1* alleles

